# Genome reconfiguration prior to mitosis shapes the generation of adaptive immunity

**DOI:** 10.1101/762757

**Authors:** Wing Fuk Chan, Hannah D. Coughlan, Jie H.S. Zhou, Christine R. Keenan, Naiara G. Bediaga, Phillip D. Hodgkin, Gordon K. Smyth, Timothy M. Johanson, Rhys S. Allan

## Abstract

During cellular differentiation chromosome conformation is altered to support the lineage-specific transcriptional programs required for cell identity. When these changes occur in relation to cell cycle, division and time is unclear. Here we followed B lymphocytes as they differentiated from a naïve, quiescent state into antibody secreting plasma cells. Unexpectedly, we found that gene-regulatory chromosome reorganization occurred prior to the first division, in late G_1_ phase and that this configuration is maintained as the cells rapidly cycle during clonal expansion. A second wave of architectural changes also occurred later as cells differentiated into plasma cells and this was associated with increased time in G_1_ phase. These data provide an explanation for how lymphocyte fate is imprinted prior to the first division and suggest that chromosome reconfiguration is spatiotemporally separated from DNA replication and mitosis to ensure the implementation of a gene regulatory program that controls the differentiation process required for the generation of immunity.

**One Sentence Summary:** Discrete waves of genome reorganization, spatiotemporally separated from DNA replication and mitosis, control cell differentiation.

## Main Text

Higher-order chromosome structure is important for packing large amounts of DNA into the nuclear space. It also plays a critical role in supporting gene expression via kilobase long DNA loops formed between gene promoters and distant regulatory elements such as enhancers, which are often housed in megabase scale topologically associated domains (TADs). Recent work has shown that these chromosome structures can be altered to drive cell type-specific gene regulatory programs(*1-3*). It is of great interest to understand how such ornate structure is established and maintained through the pressures of cell proliferation and differentiation to support cell identity. Progression through cell cycle exposes chromosomes to a number of biophysical challenges such as DNA replication and mitosis that potentially impact the architecture required for gene regulation. It has been proposed that DNA replication is a time when this architecture could be remodeled(*4-6*). Other groups have suggested that the hours after mitosis represent an opportunity for chromosome reorganization(*7-11*). However, to date, few studies have examined the dynamics of gene-level chromosome architecture across cell cycle and division when primary cells are exposed to differentiation signals.

Here we directly address this issue by studying how chromosome conformation changed to support the gene expression that drives the development of antibody-secreting B lymphocytes. Naive B cells reside in the secondary lymphoid organs in a quiescent state (G_0_ of cell cycle). During the initiation of an immune reaction they process activation signals from pathogens and immune accessory cells before entering into the proliferative response phase. The cellular events leading up to the first division take a relatively long time, compared to time spent in subsequent divisions. Moreover, within this first cell cycle, cells spend a majority of time in G_1_ phase, prior to shorter times in S phase, G_2_ phase and then mitosis, which occurs approximately 30hrs after the initial stimulus(*12, 13*). Evidence suggests that the lag prior to the first division is where T and B lymphocyte fate may be imprinted and transmitted to clonal descendants(*13-18*). Subsequently, activated B cells undergo rapid cell proliferation, dividing approximately every 8-10 hrs while spending a very short time in G_1_. This process of rapid cell proliferation is essential for the clonal expansion that underlies the antigen-specific immune response(*19*). These activated B cells differentiate in a division-linked fashion into antibody-secreting plasma cells and also form long-lived memory cells which are able to rapidly become antibody secreting cells upon rechallenge(*20, 21*). Surprisingly, little is known about how the genome is reorganized to support this differentiation process. Previously we have found that naive B cells substantially alter their genome organization as they undergo many rounds of division and become differentiated plasmablasts(*3*). To further understand when these changes arose and their association with transcriptional programs we here examined naïve B cells, B cells immediately following activation with lipopolysaccharide (three and ten hours post-activation), those just prior to the first division (33 hours, imminent division), the massively expanded B cell population (96 hours post-activation) and, finally, terminally differentiated plasmablasts (Fig 1 A & Supp Fig 1 A-E). We then used RNA-Seq to examine gene expression and *in situ* HiC to reveal three-dimensional genome structure. Transcriptomic analysis revealed that the majority of differentially expressed genes (DEs) occur prior to the first division (Fig 1 B and C, Supp Fig 1 F, Supp Table 1). As such, the expression of 5838 genes are significantly altered in the first three hours post-activation, a further 3963 in the following 7 hours and a further 1877 before the first division. A comparatively small number of transcriptional changes (1371 DEs) are observed as B cells clonally expand. A second wave of transcriptional change (3340 DEs) marks differentiation into plasmablasts. In contrast, our *in situ* HiC data showed very few genome organizational changes in the first 10 hours after activation (Fig 1D and E). In fact, we revealed that the major genome organizational changes occurred in two distinct waves; the first occurring just prior to the first cell division (10628 differential interactions; DIs), and the second upon plasmablast differentiation (6784 DIs)(Supp Table 2). The changes we observed are not made up of large-scale compartment switches (Supp Fig G) or TAD-level alterations (Supp Fig H-K) apart from in the imminent division population where the larger ratio of decreased DIs is likely driven by the loss of TAD structure known to be associated with chromosome condensation prior to mitosis(*22, 23*). The majority of the DIs we observed are made up of interactions spanning less that 1MB in distance and thus likely correspond to long-range gene promoter-enhancer interactions. For example, three-dimensional structure around the *Twistnb* gene, which encodes a component of the RNA polymerase I complex, diminishes in the first wave of organizational changes, presumably when the RNA polymerase I complex is no longer required to produce ribosomal RNAs and *Twistnb* expression is no longer required. As such, strong three-dimensional connections between the *Twistnb* promoter and long-range enhancers are detected prior to the first activation-induced division but not after (Fig 1 F). In contrast, DNA structure around the *Bcl6* gene models the second wave of organizational change, with relatively stable long-range interactions occurring both pre- and post-activation until plasmablast differentiation when this structure is lost (Fig 1 F). Interestingly, the organizational changes around the *Bcl6* gene, among others (Supp Fig 1 L), reflect its expression pattern, suggesting that chromosome structure potentially plays a role in regulating *Bcl6* expression. Interestingly, we also observed a loss of very long-range interactions (over 3MB) and a gain in shorter-range interactions after B cell activation which has been previously linked to chromosome decondensation(*24*) that peaks in expanded population (Supp Fig 1M).

**Figure 1.**
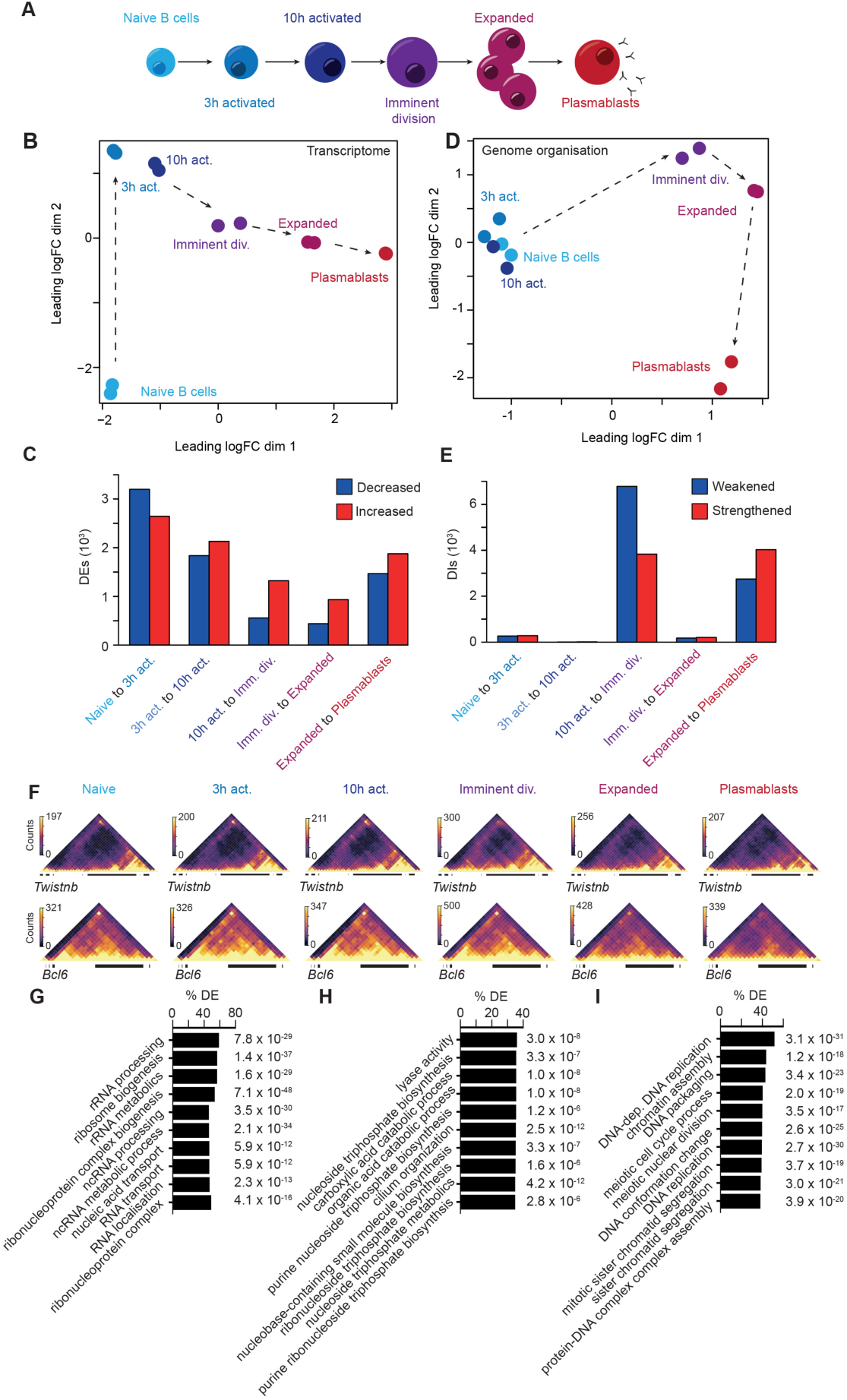
Waves of genome reorganization bookend plasmablast differentiation. **a**, Stages of B cell activation and differentiation **b**, Multidimensional-scaling plot of RNA-seq data of B cell activation **c**, Numbers of differentially expressed genes between B cell activation stages calculated with fold changes significantly above 1.5 (Treat FDR < 0.05). Blue indicates decrease in expression level with activation. Red indicates increase in expression level with activation. **d**, Multidimensional-scaling plot of B cell activation demonstrating two dominant waves of genome organisational change. **e**, Numbers of differential interactions (DIs) between B cell activation stages. Blue indicates weakening of organisation with activation. Red indicates strengthening of organisation with activation. **f**, *In-situ* Hi-C contact matrices showing genome organisation at the *Twistnb* (chr12:33.2-35.3Mb) and *Bcl6* (chr16:23.85-25.15Mb) loci at each stage of B cell activation. Color scale indicates number of reads per bin pair. **g**, Gene ontological terms with genes increasing in differential expression between the naïve and 3 hr activated B cells over-represented. Groups ranked according to percentage of groups genes that are differentially expressed. **h**, Gene ontological terms with genes increasing in differential expression between the 3 hr and 10 hr activated B cells over-represented. Groups ranked according to percentage of genes that are differentially expressed. **i**, Gene ontological terms with genes increasing in differential expression between the 10 hr activated B cells and those imminently to divide over-represented. Groups ranked according to percentage of groups genes that are differentially expressed.

In addition to identifying highly stage-restricted waves of organizational change, one of which occurs prior to the first cell division, our examination of three-dimensional structure during B cell differentiation highlights two particularly interesting points. The first is that given the absence of early activation-induced genome organizational changes, the rapid and dramatic transcriptional changes that occur immediately post-activation (Fig 1 B and C) are either driven independently of three-dimensional structure or rely on pre-existing structures. Indeed analysis of chromatin loops in naïve B cells revealed a significant link between pre-existing loop structure and subsequent gene expression changes at 3hrs after activation (*P* = 2.46 × 10^−6^)(Supp Table 3). The lack of early changes in genome organization may be explained by the transcriptional focus of the cell at these times. For example, in the first three hours post-activation gene ontological analysis of transcriptional changes (Supp Fig 1 N-O) revealed that genes involved in ribosome biogenesis are significantly enriched (Fig 1 G)(Supp Table 4), likely reflecting the B cells immediate response to activation. Between 3 and 10 hours we observed enrichment for metabolic genes (Fig 1 H)(Supp Table 4). However, the predicted function of the transcriptional changes in the hours just prior to the first cell division are almost exclusively involved in chromatin remodeling and DNA conformation change (Fig 1 I)(Supp Table 4) potentially providing an explanation as to why genome reorganization is not an early event after B cell activation. The second point of interest is the relatively small number of organizational changes that occur as a B cell clonally expands. As such, we detect few DIs between imminently dividing B cells and the massively expanded B cell population (Fig 1 D and E). This suggests that the large numbers of structures that B cells create prior to the first division are stable over the numerous subsequent divisions and would account for the striking familial symmetry in fates observed by long-term imaging(*12, 25*). This may also reflect the time restrictions imposed by rapid clonal expansion.

Our HiC data suggested that the changes in genome organization we saw were likely promoter-enhancer interactions. Linking global genome organization with gene expression has been a longstanding challenge. Therefore to link these changes to alterations in gene expression we developed a novel strategy to interrogate our organizational data. Put simply, we set a 10kb window around all protein-coding promoters and identify all DNA-DNA interactions that exist between the promoter and any other genomic region (Fig 2 A). Using the statistical framework of the R package edgeR, we determine significant differences in promoter interactivity between differentiation stages (Supp Fig 2A-D). These significantly altered promoters are known as Differentially Interacting Promoters (DIPs). Simply plotting DIPs between all six stages of B cell differentiation we find that DIPs separate the populations similarly to DEs or DIs (Fig 2 B)(Supp Table 5). Furthermore, if we examine the distribution of DIPs across all transitions we find, similar to DIs, they occur in two distinct waves; between ten hours post-activation and imminent first division and at the final transition to plasmablast (Fig 2 C, Supp Fig 2 E and F)(Supp Table 5). In addition to examining changes at each differentiation transition we group DIPs based on promoter interaction changes across the six stages of differentiation. This pattern analysis reveals 20 distinct patterns (Supp Table 6). However, the majority of DIPs can be grouped into just four patterns; increased or decreased at the imminent division population then remaining unchanged at all further transitions (2838 DIPS), or increased or decreased only upon terminally differentiating into plasmablasts (632 DIPs) (Fig 2 D and E, Supp Fig 2 G). Examining our HiC data in this way highlights the stability but also specificity of gene promoter interactions during B cell differentiation. Examining the interactivity of key B cell genes, we find interactivity reflects expression (Fig 2 F). For example, similar to their patterns of expression, the promoters of *Prdm1* and *Ell2* show increasing levels of interaction as B cell differentiation progresses. Conversely, the promoters of *Bcl6, Myc, Id2, Pax5* and *Bach2* show dramatic loss of interactions at the plasmablast stage, showing striking similarity to their patterns of expression (Fig 2 F). A broader analysis of gene ontology of DIP genes, both at each transition and within dominant patterns of change, we find that the related biological processes fit with those associated with the particular stage of differentiation (Supp Table 7 and 8). For example, promoters that showed significant and stable increases in interaction between 10 hours post-activation though prior to S phase associate with cell cycle and B cell proliferation, among others (Supp Table 7 and 8).

**Figure 2.**
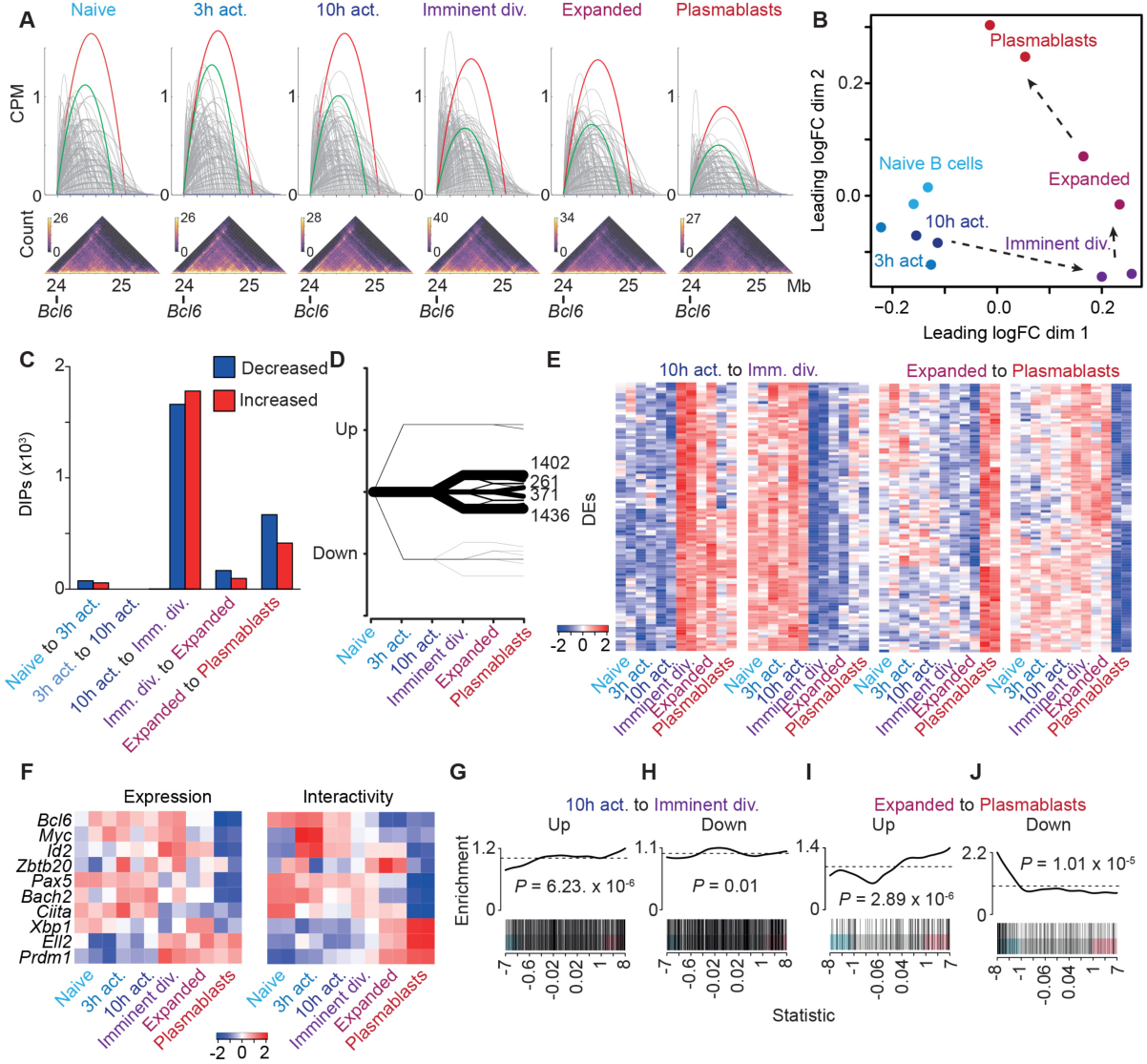
Gene promoter interactivity reveals direct link between reorganization and transcription. **a**, Schematic showing how a promoter interaction count is determined. Each interaction with the *Bcl6* promoter is shown as an arc. The sum of these interactions in counts per million (CPM) is the promoter interaction count. The red and green arcs show select interactions that represent dominant looping interactions. Below are the corresponding *in-situ* contact matrices (10kbp resolution). Color scale indicates number of reads per interaction. **b**, Multidimensional-scaling plot of the filtered and normalized logCPM values for interacting promoters during B cell activation. **c**, Bar plot showing the number of differentially interacting promoters (DIPs) between each transition of B cell activation. Blue indicates decreased interactivity. Red indicates increased interactivity. **d**, Plot showing the patterns of change and numbers of DIPs during B cell activation. Line thickness represents number of DIPs in each pattern. **e**, Heatmap of logCPM of the top 100 DIPs by false discovery rate in patterns of change in DIPs. The patterns shown represent the two dominant waves of organisational change, either exclusively increasing or decreasing at the 10 hour activation to imminent division transition or the expanded to plasmablasts transition. **f**, Heatmaps showing change in expression (logRPKM) and change in promoter interactions (logCPM) in key B cell differentiation genes across B cell activation. **g**, Barcode enrichment plot showing the correlation between promoters that significantly increase in promoter interactions as B cells prepare to divide (10 hrs activation to imminent division) with increases in gene expression at the same transition (Fig 1). Genes are ordered on the plot from right to left (x-axis) from most upregulated to most downregulated according to the moderated F-statistic. P-value was calculated with the fry test. **h**, Barcode enrichment plot showing the correlation between promoters that significantly decrease in promoter interactions as B cells prepare to divide (10 hrs activation to imminent division) with decreases in gene expression at the same transition (Fig 1). Genes are ordered on the plot from right to left (x-axis) from most upregulated to most downregulated according to the moderated F-statistic. P-value was calculated with the fry test. **i**, Barcode enrichment plot showing the correlation between promoters that significantly increase in promoter interactions as B cells differentiate into plasmablasts (expanded to plasmablast) with increases in gene expression at the same transition (Fig 1). Genes are ordered on the plot from right to left (x-axis) from most upregulated to most downregulated according to the moderated F-statistic. P-value was calculated with the fry test. **j**, Barcode enrichment plot showing the correlation between promoters that significantly decrease in promoter interactions as B cells differentiate into plasmablasts (expanded to plasmablast) with decreases in gene expression at the same transition (Fig 1). Genes are ordered on the plot from right to left (x-axis) from most upregulated to most downregulated according to the moderated F-statistic. P-value was calculated with the fry test.

While select genes appear to show a positive association between expression and organization, expanding our analysis to include all genes revealed a significant correlation between the DIPs at the two major waves of promoter interaction change and expression of the associated genes (Figure 2 G-J). For example, genes that increase expression between 10 hours post-activation but prior to the first division are significantly correlated with DIPs that also increase. The same pattern is observed in DEs during plasmablast differentiation (Fig 2 G). This suggests that the dominant function of promoter-enhancer interaction during B cell differentiation is to drive expression. Furthermore, it suggests that genome organization data alone could be used to infer gene expression in other systems.

Our data has identified that the genome architecture that defines the clonally expanded population is established prior to the first mitosis. It is currently unclear if genome reorganization occurs throughout interphase or at a specific stage of cell cycle. To address this issue we utilized the Fluorescent Ubiquitination-based Cell Cycle Indicator (FUCCI) system(*26*) to isolate imminently dividing B cells specifically in the G_1_ stage of the cell cycle (Fig 3 A and Supp Fig 3 A). While imminently dividing these cells are yet to enter the DNA synthesis phase. Using a DIPs analysis to compare this pre-S phase population to the bulk imminently dividing B cells and other stages of differentiation allows greater dissection of the contribution of cell-cycle dependent changes on the observed DIs (Fig 2 C). Remarkably, we found that although there were many DIPs between 10 hours and pre-S (Fig 3 B and C)(Supp Table 9), we observed very few between pre-S and imminent division demonstrating that genome reorganization occurs prior to DNA synthesis.

**Figure 3.**
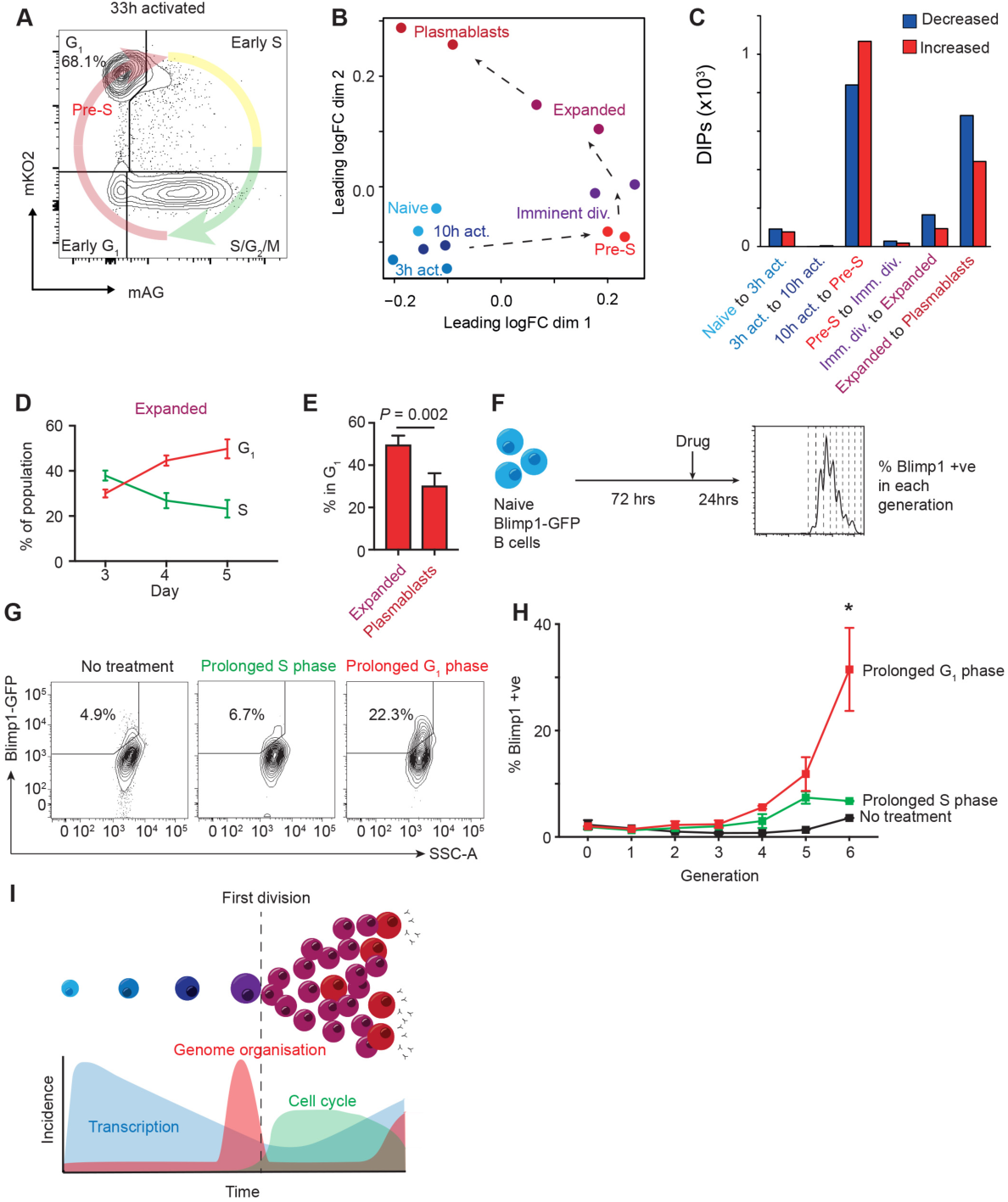
Genome organizational changes occur prior to DNA synthesis and are linked to prolonged time in G_1_. **a**, FACS plot of G_1_ stage FUCCI B cells that were FACS purified just prior to the first division (33hrs). **b**, Multidimensional-scaling plot of the filtered and normalized logCPM values for interacting promoters during B cell activation with Pre-S samples included. **c**, Bar plot showing the number of differentially interacting promoters (DIPs) between each transition of B cell activation with Pre-S samples included. Blue indicates decreased promoter interactions. Red indicates increased promoter interactions. **d**, The percentage of the expanded B cell population in the G_1_ or S phase of the cell cycle, as determined by FUCCI, at days 3, 4 and 5 of culture. **e**, The percentage of the expanded and plasmablast population in the G_1_ phase of the cell cycle, as determined by FUCCI, at day 5 of culture. The data were using an unpaired student’s *t*-test with Welch’s correction. Mean+/-SD from 4 experiments is shown. **f**, Schematic of experimental set-up for cell cycle manipulations. **g**, FACS plots showing Blimp-GFP B cells cultured with L-mimosine (prolonged S phase), Purvalonal A (prolonged G1 phase) for 24 hours or no treatment. Gate and percentage show percentage of differentiated plasmablasts (Blimp +ve) within generation 6 of the culture. **h**, Percentage of plasmablasts (Blimp +ve) within each generation of the Blimp-GFP B cell 4 day cultures when treated with Purvalonal A (prolonged G1 phase), L-mimosine (prolonged S phase) or no treatment. The data were analysed using a two-way ANOVA. Mean+/-SD from 4 experiments is shown. **i**, Model of relationship between transcription, genome organisation and cell cycle.

Given the link between G_1_ phase and genome organization we next examined whether the length of G_1_ was prolonged during the second wave of organizational change. As such, using the FUCCI system to examine the final stages of B cell differentiation we find that as B cells prepare to differentiate into plasmablasts a greater proportion are found in G_1_, consistent with spending increasing amounts of time at this stage of the cell cycle (Fig 3 D). Once differentiated, plasmablasts appear to spend significantly less time in G_1_ (Fig 3 E, Supp Fig 3 B). Thus, again we observe a naturally prolonged G_1_ phase associated with major genome organizational change. Given this association, we then tested whether artificially lengthening particular phases of the cell cycle could promote differentiation (Fig 3 F). We show that treatment of weakly differentiating, 2 day CD40/IL-4 stimulated Blimp1-GFP naïve B cell cultures with Purvalonol A [a cyclin-dependent kinase inhibitor that prolongs the G_1_ phase of the cell cycle(*27*)] drives significantly more plasmablast differentiation than treatment with L-mimosine [an L-alpha-amino acid that disrupts DNA synthesis and prolongs the S phase of the cell cycle(*28*)] or no treatment (Fig 3 G and H). These results are in line with our previous data(*29*) and supports the idea that the time spent in the G_1_ phase of the cell cycle might be limiting and particularly important for cell fate decisions, potentially to allow the numerous genome organizational changes to occur.

Over the past two decades a groundswell of evidence has demonstrated that lymphocyte fate decisions are imprinted within a day of antigen exposure, likely prior to the first division(*13-18, 30-33*). The data presented herein provides a genome-level explanation as to how this can occur. We unequivocally show that alterations in gene-regulatory chromosome conformation occur in mid-late G_1_ and are largely maintained through multiple divisions as the cells clonally expand (see to model in Fig 3I), consistent with high levels of clonal symmetry(*12, 25*). Our data also suggests that the rapid cell cycle associated with shortened G_1_ phase might be possible with little genome restructuring, enabling cells to focus on expanding prior to differentiation during a second wave of reorganization that is linked to increased time in G_1_ (Fig 3).

Our results also reveal an unappreciated spatiotemporal separation of gene-regulatory chromosome reorganization from DNA replication and mitosis. It has previously been considered that DNA synthesis provided an opportunity to remodel gene structure(*5, 34-36*) or that chromosome rewiring to support gene expression during differentiation would occur after mitosis when chromosomes decondense(*8*). Overall, we propose that genome reconfiguration is partitioned from the demanding processes of DNA replication and mitosis to ensure the safe implementation of a gene regulatory program required for the generation of cellular immunity.

## Methods

### Mice

Experiments were performed on male animals 6–12 weeks of age on a C57BL/6 Pep^3*b*^ background. Blimp-GFP and FUCCI transgenic mice were described previously (*26, 37, 38*). All mice were maintained at The Walter and Eliza Hall Institute Animal Facility under specific-pathogen-free conditions. All males were randomly chosen from the relevant pool. All experiments were approved by The Walter and Eliza Hall Institute Animal Ethics Committee and performed under the Australian code for the care and use of animals for scientific purposes. Results were analyzed without blinding of grouping.

### B cells activation and differentiation

B lymphocytes were cultured in RPMI 1640 with 2mM GlutaMAX, 50 µM β-mercaptoethanol and 10% heat-inactivated fetal calf serum (FCS). Splenocytes were obtained from male C57BL/6 mice with red cell lysis and naïve resting B cells (TCRβ-CD19+ B220+ IgM+ IgD^hi^) were sorted on BD FACS Aria III. Naïve resting B cells were activated after isolation via negative selection (Miltenyi, 130-090-862). Isolated cells were labelled with 10 µM CellTrace Violet (CTV, Invitrogen), then cultured at a density of 1×10^6^ cells/ml and stimulated with 25 µg/ml LPS (Salmonella typhosa origin, Sigma). Activated but undivided cells (TCRβ-CD19+ CD69+ CTV^highest^) were stained and sorted at the corresponding time point. FUCCI B cells were processed as above, except sorted as CD19+ CD69+ CTV^highest^ mAG-mKO2^hi^ at 33 hours post-stimulation. For long-term activation, negatively isolated B cells were cultured at a density of 1×10^5^ cells/ml and stimulated with 25µg/ml LPS. Expanded activated B cells (CD22+ CD138-) and plasmablasts (CD22-CD138+) were harvested on day 4 post-stimulation. All samples are prepared in biological duplicates for *in situ* Hi-C and RNA-seq.

### Manipulating cell cycle

B lymphocytes were stimulated with 10 µg/mL anti-CD40 (1C10) and 500 units/mL IL-4 for 72 hours at 2 x10^5^ cells/mL before being incubated with 6 µM Purvalonol A (Sigma) or 150 µM L-mimosine (Sigma) for 24 hours.

### *In situ* Hi-C

*In situ* Hi-C was performed as previously described (*39*). Libraries were sequenced on an Illumina NextSeq 500 to produce 75-bp paired-end reads. Approximately 200 million read pairs were generated per sample.

### Hi-C pre-processing

The data pre-processing and analysis was performed as previously described with changes in parameters (*3*). In brief, each sample was aligned to the mm10 genome using the *diffHic* package v1.14.0 (*40*) which utilizes cutadapt v0.9.5 (*41*) and bowtie2 v2.2.5 (*42*) for alignment. The resultant BAM file was sorted by read name, the FixMateInformation command from the Picard suite v1.117 (https://broadinstitute.github.io/picard/) was applied, duplicate reads were marked and then re-sorted by name. Read pairs were determined to be dangling ends and removed if the pairs of inward-facing reads or outward-facing reads on the same chromosome were separated by less than ∼1000 bp for inward-facing reads and ∼10000 bp for outward-facing reads. Read pairs with fragment sizes above ∼1000 bp were removed. An estimate of alignment error was obtained by comparing the mapping location of the 3’ segment of each chimeric read with that of the 5’ segment of its mate. A mapping error was determined to be present if the two segments were not inward-facing and separated by less than 1000 bp, and around 1-3% were estimated to have errors.

### Detecting differential interactions (DIs)

Differential interactions (DIs) between the different libraries were detected using the *diffHic* package v1.16.0 (*40*). Read pairs were counted into 50 kbp bin pairs for all autosomes. Bins were discarded if had a count of less than 10, contained blacklisted genomic regions as defined by ENCODE for mm10 (*43*) or were within a centromeric or telomeric region. Filtering of bin-pairs was performed using the filterDirect function, where bin pairs were only retained if they had average interaction intensities more than 4-fold higher than the background ligation frequency. The ligation frequency was estimated from the inter-chromosomal bin pairs from a 1 Mbp bin-pair count matrix. Diagonal bin pairs were also removed. The counts were normalized between libraries using a loess-based approach with bin pairs less than 1.5 Mbp from the diagonal normalized separately from other bin pairs. Tests for DIs were performed using the quasi-likelihood (QL) framework (*44, 45*) of the *edgeR* package v3.26.5 (*46*). A design matrix was constructed using a one-way layout that specified the cell group to which each library belonged. A mean-dependent trend was fitted to the negative binomial dispersions with the estimateDisp function. A generalized linear model (GLM) was fitted to the counts for each bin pair (*47*), and the QL dispersion was estimated from the GLM deviance with the glmQLFit function. The QL dispersions were then squeezed toward a second mean-dependent trend, using a robust empirical Bayes strategy (*48*). A p-value was computed against the null hypothesis for each bin pair using the QL F-test. P-values were adjusted for multiple testing using the Benjamini-Hochberg method. A DI was defined as a bin pair with a false discovery rate (FDR) below 5%. DIs adjacent in the interaction space were aggregated into clusters using the diClusters function to produce clustered DIs. DIs were merged into a cluster if they overlapped in the interaction space, to a maximum cluster size of 500 kbp. The significance threshold for each bin pair was defined such that the cluster-level FDR was controlled at 5%. Cluster statistics were computed using the *csaw* package v1.18.0 (*49*). Overlaps between unclustered bin pairs and genomic intervals were performed using the InteractionSet package(*50*).

### Detecting TAD boundaries

TAD were detected with the TADbit v0.2.0.5 python based software (*51*). Read pairs were counted into 50 kbp bin pairs (with bin boundaries rounded up to the nearest MboI restriction site) using the squareCounts function of diffHic with no filter. This yielded a count matrix containing a read pair count for each bin pair in each library. The count matrix was converted into a contact matrix for each somatic chromosome with the inflate function of the *InteractionSet* package v1.12.0 (*45*). Replicate contact matrices were summed. TADs were detected for each chromosome with the function find tad of the TADbit software. Only TADs boundaries with a score of 7 or higher were included in the results.

### Detecting differential TAD boundaries

Differential TAD boundaries between all Hi-C libraries were detected using the *diffHic* package(*40*). To identify boundaries, the directionality index (*52*) for each genomic region was determined by counting the number of read pairs mapped between the genomic region 1 Mbp upstream and separately 1Mbp downstream then taking the normalized difference of the counts. This produces a log-fold-change for each sample at each genomic region. Differential TAD boundaries were determined to be genomic regions were the size or the direction of the logFC between up and down stream counts changed significantly between conditions.

The domainDirections function was used to determine counts for 100 kbp genomic regions for 1 Mbp up- and downstream. A DGEList was constructed from the counts and library size. Low average abundance regions (calculated across all samples) of less than 1 were removed. Tests for differential TAD boundaries were performed again using the QL framework of the edgeR package as described in the Differential interaction analysis section. A log-FC threshold of log2(1.1) was applied when performing the statistical test with glmTreat from the edgeR package (*53*). A design matrix was constructed with cell type specific coefficients for the log-fold change between up- and downstream counts for each cell type. It also contains library specific blocking factor. A p-value was computed against a log_2_-fold change of 1.1 for each region using glmTreat and adjusted for multiple testing using the Benjamini-Hochberg method (FDR below 5%). A boundary was considered to be strengthened from one cell type to another if the absolute value of the log-fold change between up- and downstream counts has increased.

### Detection of A/B compartments

A/B compartments were identified across all the stages of B cell activation at a resolution of 100 kB with the method outlined by Lieberman-Aiden et al.(*54*) with the HOMER HiC pipeline (*55*). After processing with the diffHic pipeline, libraries were converted to the HiC summary format with R. Then input-tag directions were created for each library with the makeTagDirectory function, with the genome (mm10) and restriction-enzyme-recognition site (GATC) specified. Biological-replicate tag directories for each cell type were summed. The runHiCpca.pl function was used on each library with -res 100000 and the genome (mm10) specified to perform principal component analysis to identify compartments. To identify changes in A/B compartments between libraries, the getHiCcorrDiff.pl function was used to directly calculate the difference in correlation profiles.

### Detecting differentially interacting promoters (DIPs)

Differentially interacting promoters were detected using across all libraries with the diffHic package(*40*). Gene promoters were defined with the genes from the TxDb.Mmusculus.UCSC.mm10.knownGene package v3.4.7 and by applying the promoters function from the GenomicFeatures package v1.36.2 with upstream=5kbp and downstream=5kbp. Interactions between the promoters and the entire genome was counted with the connectCounts function from diffHic with filter set to 0 and the second.region=10kbp. Interchromosomal interactions were excluded along with interactions contained in blacklisted regions as defined by ENCODE, centromeres or telomeres. Interactions were filtered by abundance as a function of distance between the anchors. Using the loessFit function from the limma package v3.40.2 (*56*) with span 0.05, a loess curve was fitted to the average log counts per million (logCPM) for all libraries (calculated with the cpmByGroup function of EdgeR with log=TRUE) as a function of distance^0.25. An interaction was then required to have an abundance larger than the fitted curved plus two times the mean of the absolute values of the residuals from the loess fit. For each unique promoter, interactions were then aggregated to produce a count matrix. Low-abundance promoters were filtered using edgeR’s filterByExpr function with min.count=200 and min.totals=200. Obsolute Entrez Gene Ids were removed, as were mitochondrial, ribosomal (rRNA/ncRNA), Riken, olfractory and X or Y chromosome genes. The counts were normalized between libraries using a loess-based approach with the normOffsets function from the csaw package v1.18.0.

Two different DIPs analyses were performed; with and without the FUCCI B cells Hi-C libraries. DIPs were detected with the quasi-likelihood (QL) framework (*44, 45*) of the edgeR package. A design matrix was constructed with a one-way layout that specified the cell type. Using the promoter counts, the estimateDisp function was used to maximise the negative binomial likelihood to estimate the common dispersion across all promoters with trend=none and robust=TRUE (*57*). A generalized linear model (GLM) was fitted to the counts (*47*) and the QL dispersion was estimated from the GLM deviance with the glmQLFit function with robust=TRUE and trend.method=none. The QL dispersions were then squeezed toward a second mean-dependent trend, with a robust empirical Bayes strategy (*48*) to share information between genes. A *P-*value was calculated for each promoter using a moderated t-test with glmQLFTest. The Benjamini–Hochberg method was used to control the false discovery rate (FDR) below 5%. Heatmaps of the filtered and normalized logCPM value were plotted with the coolmap function from the limma package. Mean-difference plots (MD plots) were plotted with the plotMD function.

For each DIP in the analysis, the pattern of the promoter interactions was defined across all the transitions as either: 1 (increasing at a transition), -1 (decreasing at a transition) or 0 (not significantly changed). An individual DIP was determined to be in a pattern if it had FDR below 5% at that transition while the sign of the logFC gave the direction.

### Detecting looping interactions

Looping interactions were detected in the naive B cell libraries with a method similar to that described by Rao et al. (*39*) and as described previously (*3*) except with a bin size of 20 kbp. Established loops were defined as those with enrichment values above 0.5, with an average count across libraries greater than 5, and that were more than 60 kbp away from the diagonal. Loops were annotated with genes in the anchors with annotatePairs function from the diffHic package. The significance of the enrichment of DE genes between naïve B cells and 3 hours after activation (without logFC threshold) in the loops was performed with a pearson’s chi-squared test (df=2). The test was between the portions of genes within/not within loop anchors that are: DE genes increasing, DE genes decreasing or non-DE genes.

### Visualisation of Hi-C

Multidimensional-scaling plots were constructed with the plotMDS function in the limma package applied to the filtered and normalized logCPM values of each bin pair or promoter for each library. The distance between each pair of samples was the ‘leading log fold change’, defined as the root-mean-square average of the 500 largest log_2_ fold changes between that pair of samples.

Plaid plots were constructed using the contact matrices and the plotHic function from the Sushi R package v1.22.0 (*58*). The color palette was inferno from the viridisLite package v0.3.0 (*59*) and the range of color intensities in each plot was scaled according to the library size of the sample. The plotBedpe function of the Sushi package was used to plot the unclustered DIs as arcs where the z-score shown on the vertical access was calculated as - log_10_(*p*-value).

### RNA-seq

RNA was extracted with NucleoSpin RNA Plus (Macherey-Nagel) and subsequently quantified in a TapeStation 2200 using RNA ScreenTape (Agilent). Libraries were prepared with a TruSeq RNA sample-preparation kit (Illumina) from 500 ng RNA, as per the manufacturers’ instructions. Libraries were then amplified with KAPA HiFi HotStart ReadyMix (Kapa Biosystems) and 200 to 400 bp products were size-selected and cleaned up with AMPure XP magnetic beads (Beckman). Final libraries were quantified with TapeStation 2200 using D1000 ScreenTape for sequencing on the Illumina NextSeq 500 platform to produce 75 bp paired-end reads. Around 25 million read pairs were generated per sample.

Reads were aligned to the mm10 genome with Rsubread package v1.28.0 Genewise counts were obtained with featureCounts and Rsubread’s inbuilt Entrez gene annotation. Low-abundance genes were filtered out with the filterByExpr function. Obsolete Entrez Gene Ids were removed, as were mitochondrial, ribosomal, X or Y chromosome genes and variable immunoglobulin gene segments. Normalization was performed with the trimmed mean of M-values (TMM) method (*60*).

The differential expression analysis was conducted using the quasi-likelihood (QL) framework (*44, 45*) of edgeR. The empirical Bayes procedure of glmQLFTest was run in robust mode to increase power and protect against hypervariable genes (*48*). Differential expression was evaluated using edgeR’s glmTreat function with log-fold-change threshold set to 1.5 (*53*). This method prioritizes genes with biological significantly changes by ensuring that all differentially expressed genes have observed fold-changes significantly greater than 1.5. The Benjamini–Hochberg method was used to control the FDR below 5%. Log2 RPKM (reads per kilobase per million) were computed with edgeR’s rpkm function. MDS plots, MD plots and heatmaps were created using the plotMDS, plotMD and coolmap functions. Distances on the MDS plots correspond to leading log2-fold-changes from the top 500 differentially expressed genes.

Patterns of differential expression were determined for genes with the same method as the DIPs.

### Gene set enrichment

Tests for over-representation of gene ontology terms was performed with the goana function from the limma package. Gene set enrichment was tested using limma’s fry function and visualised using limma’s barcode enrichment plot. Terms containing less than 100 genes were removed. Categories were ranked according the percentage of genes within the category that are differentially expressed. Terms in the “CC” ontology category were also removed.

## Data accessibility

Sequence data that support the findings of this study are tabulated in the supplementary tables and are available in the GEO database under accession number GSE99151.

---

**Supplementary Figure 1.**
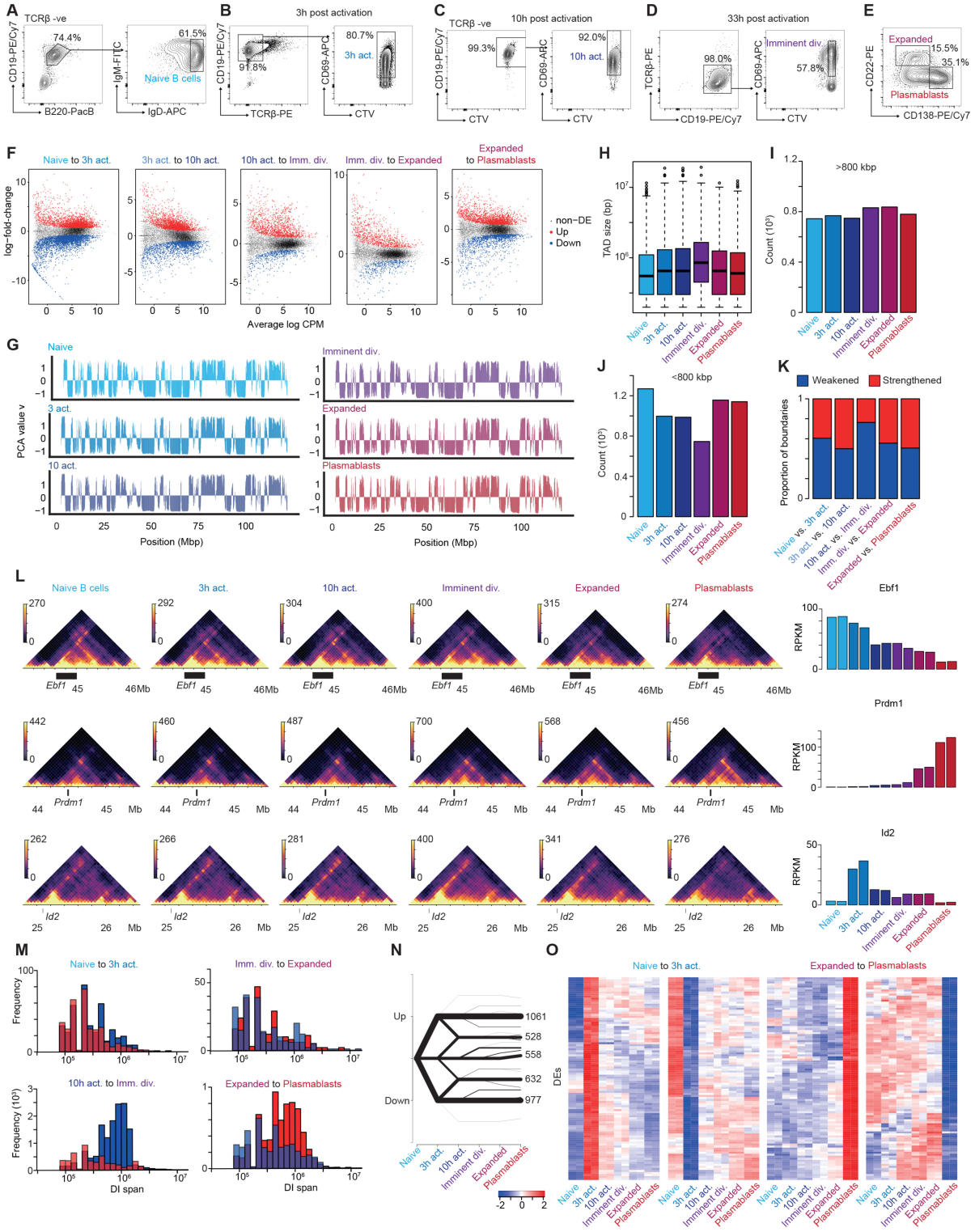
**a**, FACS plots showing strategy used to isolate naïve B cells from mouse spleen. **b**, FACS plots showing strategy used to isolate activated B cells from culture 3 hours after their activation with LPS. **c**, FACS plots showing strategy used to isolate activated B cells from culture 10 hours after their activation with LPS. **d**, FACS plots showing strategy used to isolate activated B cells from culture 33 hours after their activation with LPS. **e**, FACS plot showing strategy used to isolate the expanded B cell population and plasmablasts from culture 4 days after their activation with LPS. **f**, Mean-difference (MD) plots with differentially expressed (DE) genes highlighted in red (up) and blue (down) for each transition of B cell activation. DE genes have fold-changes significantly greater than 1.5 (Treat FDR < 0.05). Plotted on the y-axis is log-intensity ratios of the genes and plotted on the x-axis is the log-intensity averages. **g**, A/B compartmental interaction plots of *in situ* Hi-C data of the entirety of chromosome 12 at different B cell activation stages. 100 kB resolution. **h**, Distributions of TAD size (bp) from summed libraries called with TADbit at each stage of B cell activation. **i**, Number of TADs detected that are smaller than 800 kbp at each stage of B cell activation. **j**, Number of TADs detected that are larger than 800 kbp at each stage of B cell activation. **k**, Proportion of TAD boundaries strengthening or weakening at each transition of B cell activation determined by *diffHic*. **l**, *In-situ* Hi-C contact matrices showing genome organisation at the Ebf1 locus (chr4:44.2-45.5Mbp), *Prdm1* locus (chr10:43.8-45.5Mbp) and the Id2 locus (chr12: 24.8-26.5Mbp) during all stages of B cell activation, with corresponding expression changes (far right). Color scale in the contact matrices indicates number of reads per bin pair. **m**, Histogram of unclustered DI span (distance between boundaries of DNA structure) for transitions during B cell activation (50 kbp resolution). Blue indicates DIs that are decreasing in fold-change. Red indicates DIs that are increasing in fold-change. **n**, Plot showing the patterns and numbers within each pattern of DEs during B cell activation. **o**, Heatmap of logRPKMs of the top 100 DEs by false discovery rate in four select patterns expression change during B cell activation. The left panels show DEs that transiently increase or decrease in expression 3 hours post-activation before returning to their original level. The right panels show DEs that upregulate or downregulate exclusively during plasmablast differentiation.

**Supplementary Figure 2.**
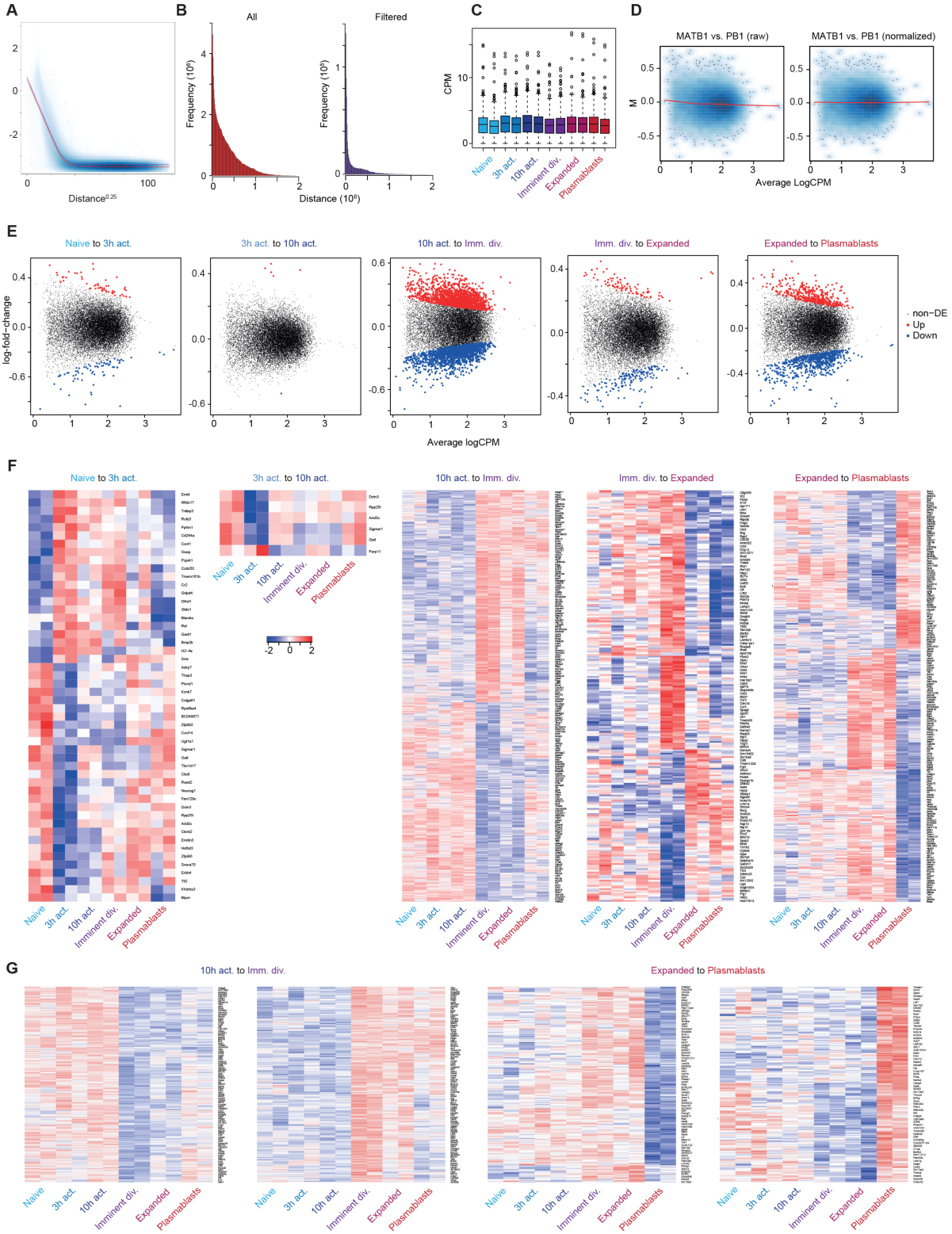
**a**, A plot of the average log counts per million (logCPM) of interactions for all libraries as a function of interaction distance to the power of 0.25. Interaction distance is the number of bp between the boundaries of an interaction. A loess curve was fitted to the data with a span of 0.05 shown in red. For an interaction to be retained it was required to exceed the fitted curved plus two times the mean of the absolute values of the residuals from the loess fit shown in purple. **b**, Histograms of the interaction distance of all interactions before and after applying filtering. **c**, Boxplots of the CPMs of the aggregated promoter counts for each library. **d**, Mean-abundance (MA) plot of a naïve B library versus a plasmablast library before and after normalization. Plotted on the y-axis is the log-fold change of the interacting promoter counts between libraries and plotted on the x-axis is the log-intensity averages. **e**, Mean-difference (MD) plots with differentially interacting promoters (DIPs) highlighted in red (up) and blue (down) for each transition of B cell activation. **f**, Heatmap of logCPMs of all DIPs at each transition of B cell activation. **g**, Heatmap of logCPMs of all DIPs in patterns of DIPs that exclusively increase or decrease at the 10 hours activation to imminent division transition and expanded to plasmablasts transition.

**Supplementary Figure 3.**
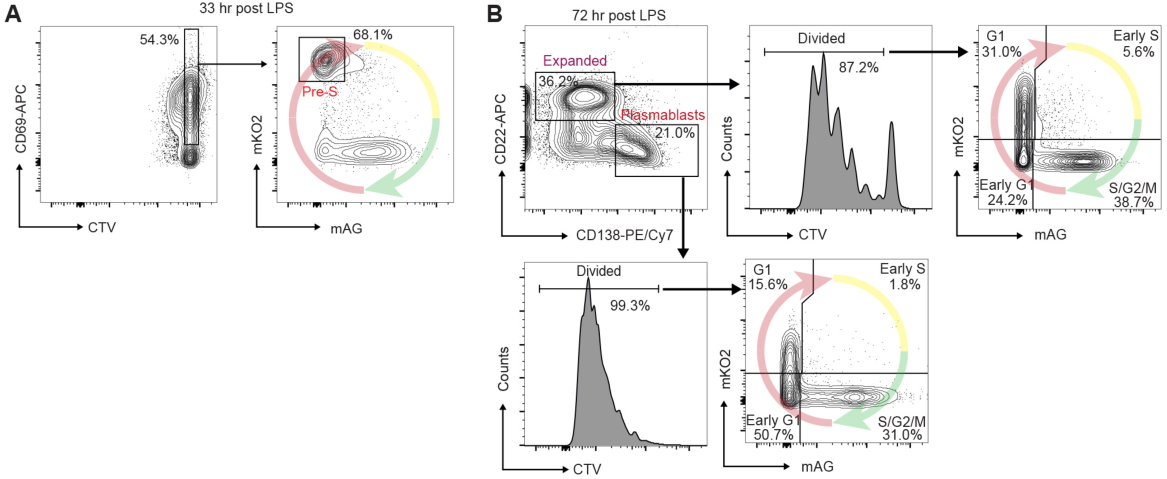
**a**, FACS plots showing strategy for isolating undivided, activated, G_1_ cells 33 hours after LPS exposure. **b**, FACS plots showing strategy for determining the cell cycle states of expanded B cells and plasmablasts in 3 day cultures.

